# Condition dependent strategies of egg size variation in the Common Eider *Somateria mollissima*

**DOI:** 10.1101/863654

**Authors:** Thomas Kjær Christensen, Thorsten Johannes Skovbjerg Balsby

**Author notes:** Corresponding author (TKC).

## Abstract

We analysed intraclutch egg-size variation in relation to clutch size and to female body condition in the Common Eider *Somateria mollissima* during an 8-year period. The aim was to assess if eiders adaptively adjusted egg size within the laying sequence in response to different clutch sizes, which potentially could optimise reproductive success through a size advantage in hatchlings. The analyses were performed on both population and individual level using data from recaptured females that changed clutch size between seasons. Based on 1,099 clutches from 812 individual females, population clutch size averaged 4.13 eggs (range: 1-6), with marked annual variation in the dominance of 4- and 5-egg clutches, which constituted *c.*70% of all clutches. Clutch size was positively related to female pre-laying body condition at both the population and individual level. Egg size varied significantly within and between clutch sizes and changes were significantly related to the laying sequence. First eggs were significantly larger in 4-egg clutches and second eggs smaller (marginally insignificant) than in 5-egg clutches, a pattern also found among individual females changing clutch size between seasons. The relationship between female pre-laying body condition, clutch size and the intraclutch egg-size pattern indicates that both clutch size and egg size is actively adapted to the pre-breeding body condition of the female. We suggest that the observed pattern of intraclutch egg-size variation reflects a finely tuned conditional dependent mechanisms that enable females in a suboptimal condition to optimize reproductive output.

## Introduction

Intraclutch egg-size variation has frequently been described in both altricial and precocial avian species [1–3]. In large long-lived precocial waterfowl species, such as geese and eiders, egg-size variation over the laying sequence generally follow a rather fixed curvilinear pattern, with increasing size from first to second and/or third egg and then a declining size over the following eggs in the laying sequence, with the final egg being the smallest [4–10]. With few exceptions, this pattern of intraclutch egg-size variation is consistently found among the most commonly laid clutch sizes within populations, occurring almost independently of clutch size, female age and condition, and time of laying [3–4, 7, 11–12]. Parsons [13] suggested that the small size of first egg was related to a lower physiological efficiency in females at the up-start of egg laying, while other studies support a physiological and hormonal, genetically based, explanation to the overall pattern of intraclutch egg-size variation [see 4]. Alternative, but not mutually exclusive hypotheses, have related intraclutch egg-size variation to the depletion of resources in laying females with egg number, incubation strategy, facilitation of hatching synchrony and differential investment in eggs with highest probability of hatching [8, 10–11, 14–22]. Despite the long-term focus on intraclutch egg-size variation, neither adaptive nor non-adaptive explanations have proved as full explanations to the observed egg size patterns.

Even though egg size in birds generally show high repeatability and heritability, and consequently is less flexible in individual birds than e.g., clutch size [cf. 23–25], egg size and intraclutch egg-size variation may have important implications for the reproductive success of individual females. In geese and eiders, laying small clutches of 4 to 6 (1-7) relatively large eggs, larger eggs within the clutch produce larger hatchlings that have a faster growth rate, higher pre-fledging survival and even a higher recruitment rate [3, 26–31, but see 32]. However, despite the apparent benefit of making adaptive changes (e.g. laying larger eggs when laying a reduced clutch size or vice versa, or changing egg size according to female conditional state or in relation to environmental conditions), such adaptive changes have not been substantially documented [see 5, 33–35]. This is surprising, as even small proportional changes in egg size that would balance egg size within clutches, is expected not to be costly to females already being physiologically upgraded for breeding [cf. 11, 24]. Likewise, avian life-history characteristics suggests that adaptive patterns in both clutch and egg size should be more prevalent in species that store resources for reproduction in the pre-breeding period (capital breeders) compared to species depending on resources available during the laying and chick-rearing period (income breeders) [36]. Thus the apparent lack of flexibility in egg size even in capital breeding birds, suggests that adaptations to varying conditional states of females, is mainly adjusted through changes in clutch size [cf. 24]. Indeed, many studies have shown that females in good condition lay larger clutches that are reproductively more successful than smaller clutches [e.g., 14, 23, 37–39].

In a long-term study of breeding eiders in Denmark, recordings of egg size showed that intraclutch egg-size variation over the laying sequence followed the common curvilinear pattern as generally described in eiders [see 7, 9]. However, we also observed that egg size in the most commonly laid clutches of 4 and 5 eggs showed a systematic difference with respect to the size of the first laid egg. Examining previous studies of eider clutch and egg size, we found a similar difference in other breeding populations, giving the impression that eiders adaptively changed the size of first laid egg depending on clutch size.

In the present study, we use data from a long-term survey of breeding eiders to test if the common eider make adaptive changes in egg size within the laying sequence dependent on clutch size. Since clutch size in the eider is closely related to female body condition, we also included female body condition in the analyses. We analyse data on a population level, as well as an individual level, based on ringed and recaptured individual females that changed clutch size within our study period of 8 years.

## Materials and Methods

### Data collection

Data on clutch size and egg size were collected annually during 1993-2000 at the island of Saltholm, Denmark (55°55’N 12°46’E). The island, covering 13.5 km^2^, holds an estimated breeding population of 4,000 to 6,000 eiders [40]. In all years, seven 45 m wide coast-to-coast transects, covering an area of c. 0.75 km^2^, were searched thoroughly for nests towards the end of the incubation period in early May. Transects were distributed evenly 1 km apart following east-west oriented UTM grid lines. Attempts were made to capture all encountered incubating females, but only data from clutches where females were caught (assumed incubating on a fully laid clutch) were included in the present analyses. Two females sitting on clutches of 7 eggs were omitted as egg size and coloration strongly indicated egg dumping in both nests. We consider that the extremely low nest densities (average 0.65 nest per 45 × 45m sampling unit ∼ 3.2 nest per ha [40]) relative to other studies generally precludes nest parasitism, which increases with nest density, being reported in colonies above 10-15 nest per ha [cf. 41]. However, we cannot reject that a few incomplete or parasitized nests, as well as some partially predated nests (in cases no egg-shell remains was found that disqualified the record) may unintentionally have been included in the present data, and hence have induced some uncontrolled variation.

During 1993 to 2000, 1,099 incubating females were captured on nests (between 79 and 174 per year), ringed with standard steel leg bands and body mass (to nearest 5 gram) and tarsus length (to nearest 1 mm) recorded. The sample included 812 individual females, of which 610 were caught once, 141 twice, 44 thrice, 12 four times, 3 five times and 2 six times.

Captures and ringing of eiders were conducted according to the practical and ethical guidelines for bird ringing and carried out under the institutional ringing license (no. 600) issued by the Ringing Centre at the Natural History Museum of Denmark.

The 1,099 clutches contained a total of 4,531 eggs. In all clutches, individual eggs were related to laying sequence (first to sixth egg) based on individual degree of staining [4, 8, 34, see also 42], a method previously applied to eiders [23]. Egg length and width were measured to nearest 0.1 mm using callipers.

Egg volume was subsequently calculated according to the formula: Volume = length × π × bredth^2^ × 0.00164 cm^3^ [9, 43]. In each year, female body condition (mass corrected for structural size) prior to egg laying (W_start_) was estimated for each individual female based on the year specific difference between body mass at capture (W_capt_) compared to average year specific body mass at hatching (W_hatch_) for a given structural size (tarsus length). We assumed a weight loss of 20 gram per day during incubation and an incubation period of 28 days [cf. 44]. Hence, individual body condition at start of egg-laying was calculated as

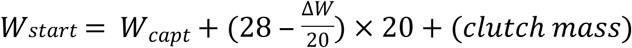

where Δw is the difference between body mass at capture and the annual average body mass at hatching for a given structural size. Annual average body mass at hatching (W_hatch_) was obtained from regression of body mass on the tarsus length (cubed) of all females captured with ducklings or hatching eggs in their nests, accounting for annual differences in average population body condition. Clutch mass was estimated setting average egg mass to 100 gram [cf. 45, 46].

To assure that clutch size differences were not responsible for differences in W_start_-estimates, we tested relations between clutch sizes and observed W_hatch_ for clutch sizes of 3 to 5 eggs in which hatching (ducklings or piping/cracking eggs) were recorded (N = 150). We used a mixed model with clutch size, tarsus^3^ and their interaction as fixed variables and year as random variable. Clutch size and the interaction clutch size * tarsus^3^ did not affect W_hatch_ (Mixed model, clutch size: F_2, 137_ = 0.23, p = 0.794; clutch size * tarsus^3^: F_2, 137_ = 0.16, p = 0.852), whereas tarsus^3^ showed a significant relation (F_1, 137_ = 11.1, p = 0.001). This analysis included an additional 112 females caught with ducklings in the nest, which were not part of the egg-size sample.

In the analyses of intraclutch egg-size variation, we only included clutches of 2 to 6 eggs (98.5% of all clutches recorded) given the uncertainty associated with clutches of 1 (likely incomplete) and 7 eggs (potentially parasitized). When analysing individual egg-laying patterns in recaptured individuals, we only included 4 and 5-egg clutches, as these were numerically dominating (> 70% in all years).

### Statistics

We tested egg size differences in first, second and third egg between clutch sizes, using a mixed model including clutch size as fixed and year as a random factor. We tested the pattern of egg size differences over the laying order among clutch sizes using a repeated measure ANOVA, to detect if the pattern with laying order differed between different clutch sizes. The model included laying order, clutch size and year, and the interaction between laying order and clutch size, and interaction between year and laying order. The model used individual nests as the repeated measure. Note that the relationship between laying order and egg size only could be compared for the number of eggs that corresponds to the smallest clutch size selected (e.g., would only test the pattern for the first four eggs in clutches with at least four eggs). We therefore conducted repeated measures ANOVA for each number of egg.

For recaptured females, the likelihood of changing her reproductive strategy with regard to clutch size in a given season was tested by relating inter-seasonal changes in clutch size to inter-seasonal changes in body mass at the start of egg laying. We classified female reproductive strategy as either reducing, maintaining or increasing clutch size. To account for multiple contributions by some individuals (up to 4 recaptures) we included individual as a random factor in the mixed model. The likelihood of a change (increasing, maintaining or reducing) in clutch size as a response to differences in pre-laying body mass were tested by a generalized linear model with a multinomial distribution. For post hoc pairwise analyses of egg volume among all clutch sizes, we used least square mean differences. The proportion of significant pair-wise differences by far exceeded 5% expected by random, and hence did not require adjustment of significance threshold. We used SAS 9.3 (SAS Institute, Cary, NC) for all analyses using Proc Mixed, Proc GLM, Proc Univariate, and Proc Freq.

## Results

### Clutch size

Average clutch size during 1993-2000 was 4.13 ± 0.03 SE (N = 1,099). Clutch size approached statistically significant variation between years, with the largest clutch sizes recorded in 1993 and 1996 and the smallest clutch size in 1997 (range: 3.97-4.30; General linear model: F_7,1091_ = 1.99, p = 0.054, R^2^ = 0.013)(Fig. 1) (S1 Table).

**Fig 1.**
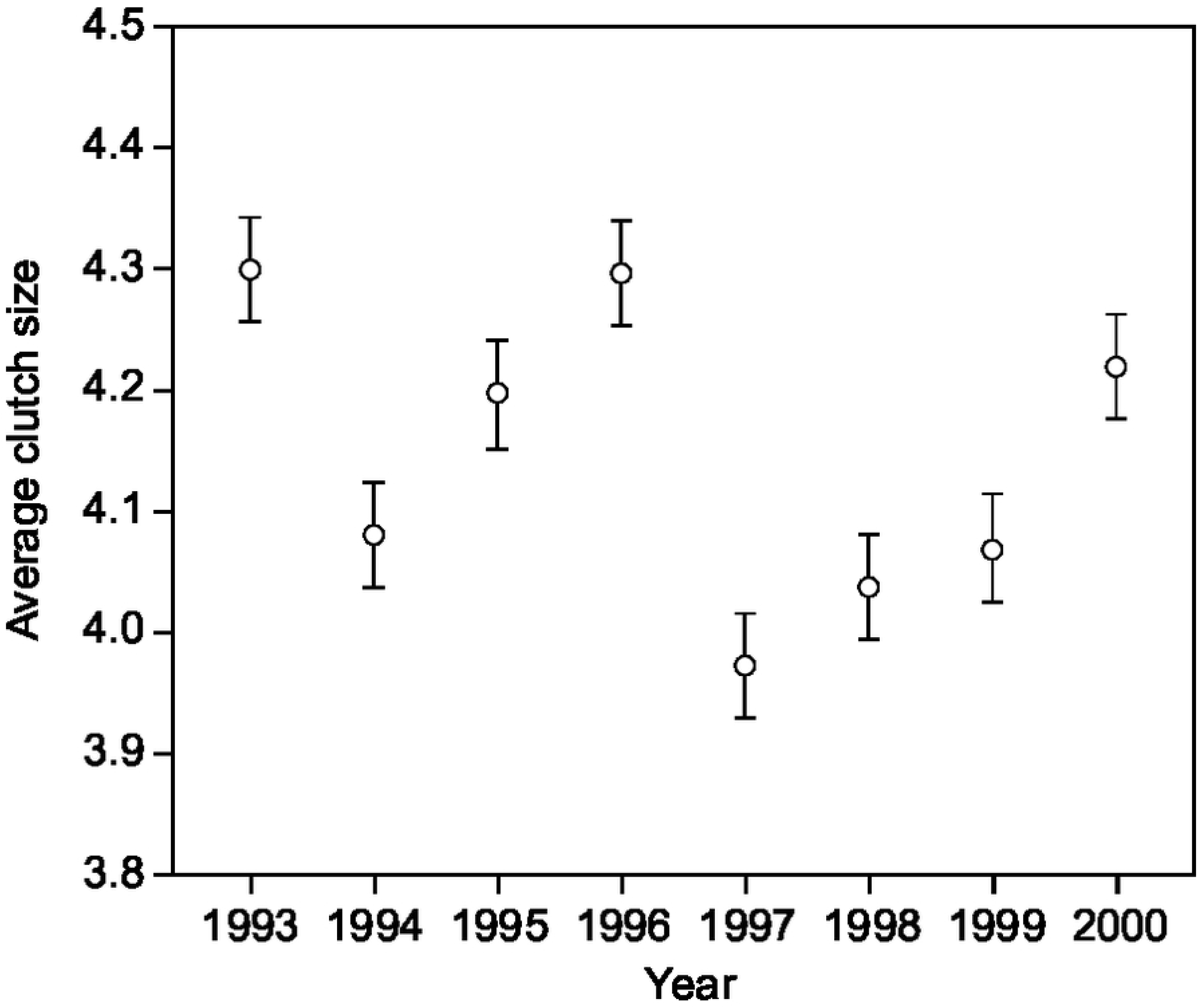
Average clutch size (±SE: standard error) for 1,099 clutches of eiders captured during incubation on the island of Saltholm during 1993-2000.

Clutches of four and 5 eggs were most frequent (annual average 35.9%, range: 24.5-47.3% and 37.8% range: 30.3-50.4% respectively). A significant (Chi-square test: χ_6_^2^ = 20.2, p < 0.005) shift in dominance of 4 and 5 egg clutches between years, affected overall population clutch size. Total clutch volume of 4-egg clutches was on average 80.3% of the volume of 5-egg clutches.

Clutch size were significantly and positively related to female body condition at the start of egg-laying (General linear model: clutch size: F_1, 809_ = 1382,53, p < 0,0001slope = 0.006).

### Clutch size for recaptured females

Of the recaptured females that reduced clutch size in a following year, the majority showed a decrease in body mass (72 had lost weight and 18 had gained weight). Of the recaptured females that increased clutch size between years, the majority showed an increase in body mass (86 had gained weight and 12 had lost weight). Among the 108 females, that did not change clutch size between years there was no distinct pattern with regard to changes in body mass (43 females lost weight and 65 females gained weight). The changes in individual female body mass at start of incubation between years significantly affected changes in clutch size (increasing, maintaining or reducing clutch size)(Generalized linear mixed model F_1, 287_ = 104.6, p > 0.0001, Estimate = 0.011). Hence, when females lost body mass relative to the previous breeding record, they were significantly more likely to reduce clutch size, as indicated by the model estimates, and vice versa.

### Intraclutch egg-size variation

Overall mean egg volume (hereafter egg size) in all clutches (N = 4,531) was 103.1 ± 0.1 cm^3^. Excluding extreme outliers of one dwarf (20.1 cm^3^) and one extremely large egg (176.7 cm^3^), egg size range was 79.4-159.5 cm^3^ (n = 4,529) and the smallest egg was 49% the size of the largest egg in the population of measurements.

Egg size showed a consistent pattern with laying order across different clutch sizes (Fig. 2). The size of the first, second and third egg differed significantly between clutch sizes (Table 1). The difference in the size of the first egg was related to a significantly larger egg in 4-egg clutches compared to all other clutch sizes (Fig. 2, S2 Table). The second egg in the laying sequence was significantly larger in 5-egg clutches compared to 2- and 3-egg clutches, almost significantly larger (p = 0.054) than in 4-egg clutches, but did not differ from the second egg in 6-egg clutches (Fig. 2) (S2 Table). The size of the third egg in 4-egg clutches was significantly larger than in 3-egg clutches, but significantly smaller than in 5- and 6-egg clutches.

**Fig 2.**
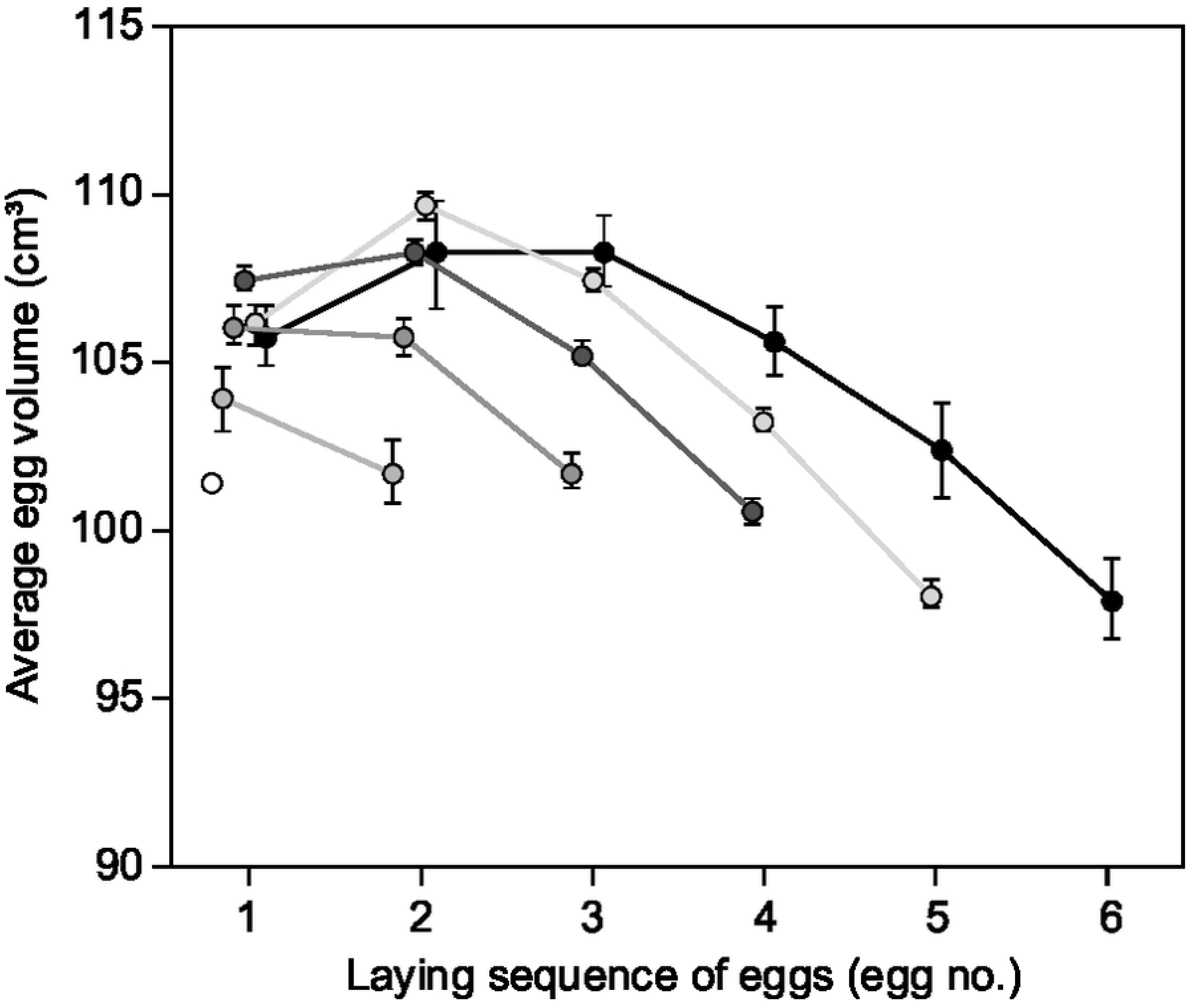
Average egg volume (±SE: standard error)) in eider clutches in relation to position in the laying sequence for females captured on nests with 1 to 6 eggs.

**Table 1.** Test of differences in volume of first, second and third eggs across clutches of 2-6 eggs. Data was analyzed by a mixed model with year as a random factor and only included first records of recaptured females.

| Volume | df | F | p |
| --- | --- | --- | --- |
| First egg | 5, 804 | 5.49 | <0.001 |
| Second egg | 4, 796 | 13.87 | <0.001 |
| Third egg | 3, 752 | 17.22 | <0.001 |

### Patterns of egg size in relation to laying order

Egg size was significantly affected by laying order in clutch sizes of 2-6 eggs as indicated by the significant effect of laying order (Table 2). More importantly, clutch size significantly affected the size of eggs within the laying sequence as indicated by the significant interactions between laying order and clutch size for clutch sizes between 2 and 6 eggs (Table 2).

**Table 2.**
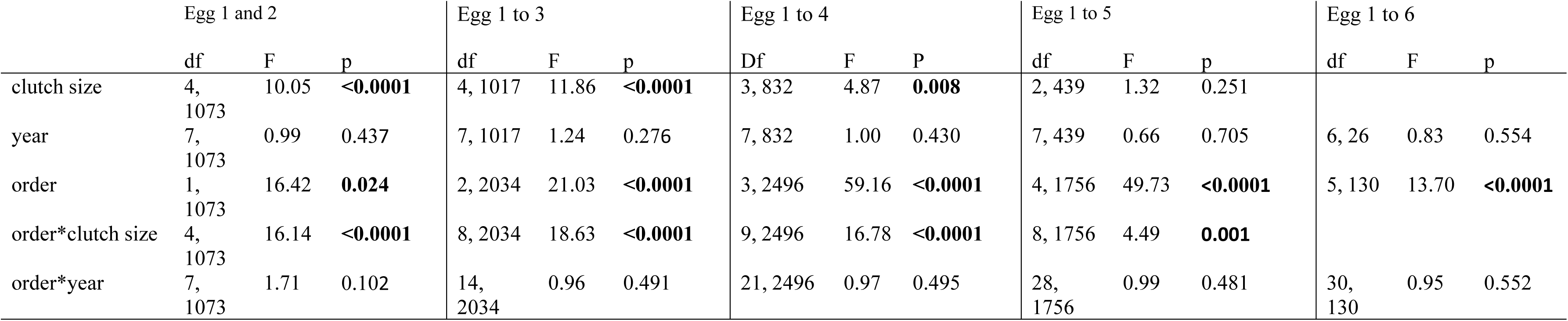
Test (repeated measures ANOVA) of egg volume changes with clutch size, year and laying order, and the interaction terms for laying order for a stepwise increase in clutch size (ex: Egg 1 and 2 includes all clutch sizes larger than one egg (2 to 6 egg clutches); Egg 1 to 5 includes clutches with 5 or more eggs (5 and 6 egg clutches)).

### Egg laying patterns within individuals

The overall population difference in intraclutch egg-size pattern of females laying 4 and 5 egg clutches, was also found in recaptured females that changed their clutch size from 4 to 5 eggs (N = 35)(Fig. 3A) and from 5 to 4 eggs (N = 46)(Fig. 3B). When laying a 4-egg clutch, the volume of first and second eggs were comparable in size (ANOVA: F_1, 68_ = 3.98, p = 0.926), but there was a marked size difference when subsequently laying a 5-egg clutch (ANOVA: F_1, 68_ = 3.98, p = 0.070). For females changing clutch size from 5 to 4 eggs, the opposite change in the size of the first and second egg was evident (ANOVA: 5-egg clutch: F_1, 90_ = 3.95, p = 0.002; 4-egg clutch: F_1, 90_ = 3.95, p = 0.875). Hence, individual females laying a 4-egg clutch increased the size of the first egg while decreasing the relative size of the second egg compared to years when laying a 5-egg clutch and vice versa. A similar, but less clear, trend was evident among females making two changes, either 4-5-4 eggs or 5-4-5 eggs, although the samples sizes were very small (N = 5 and 6 respectively) and hence subject to differences between individuals (Fig. 4).

**Fig 3.**
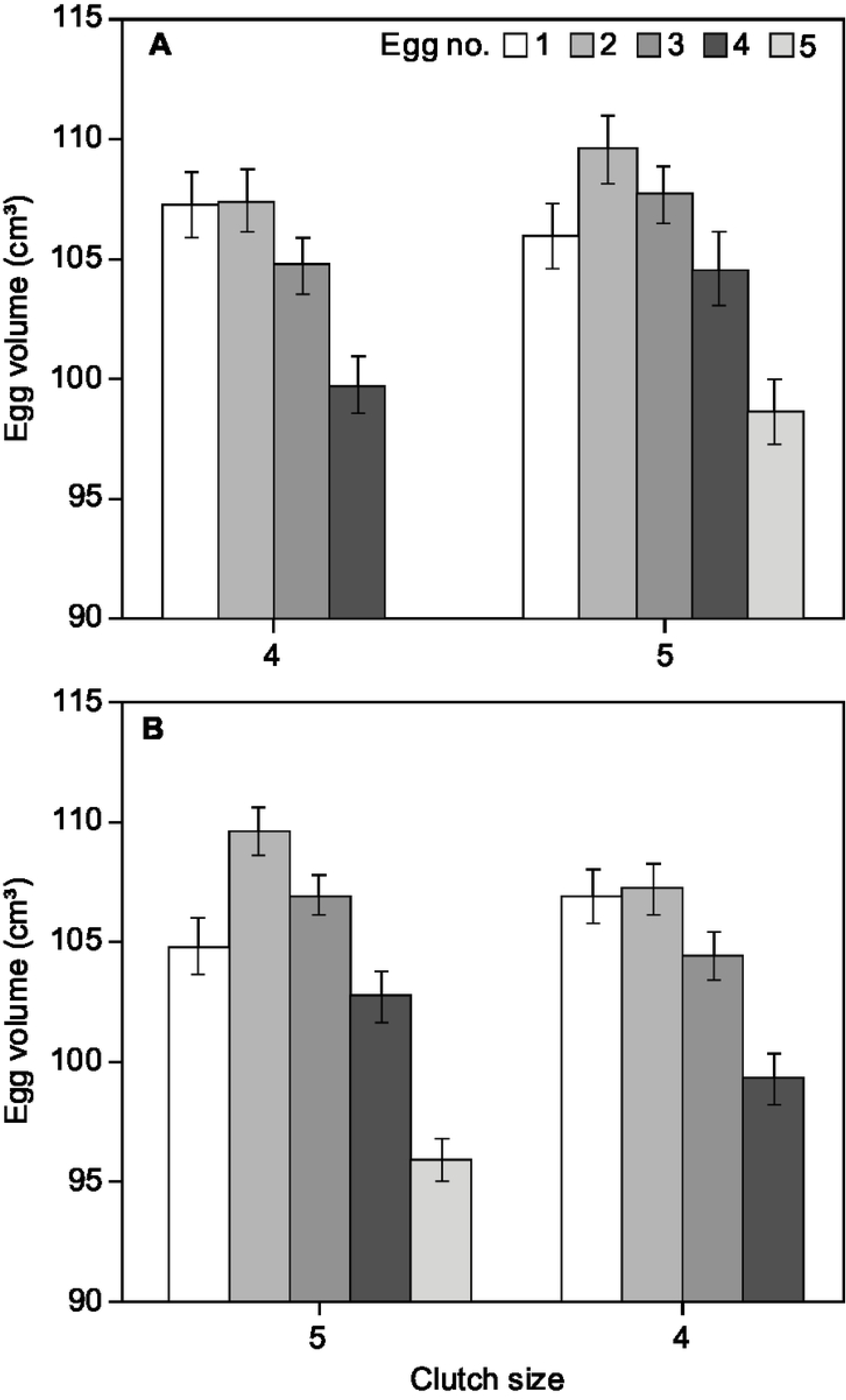
Average egg volume (±SE: standard error) for consecutive eggs in the laying sequence for individual female eiders that changed clutch size from A) 4 to 5 eggs (N = 35) and B) 5 to 4 eggs (N = 46).

**Fig 4.**
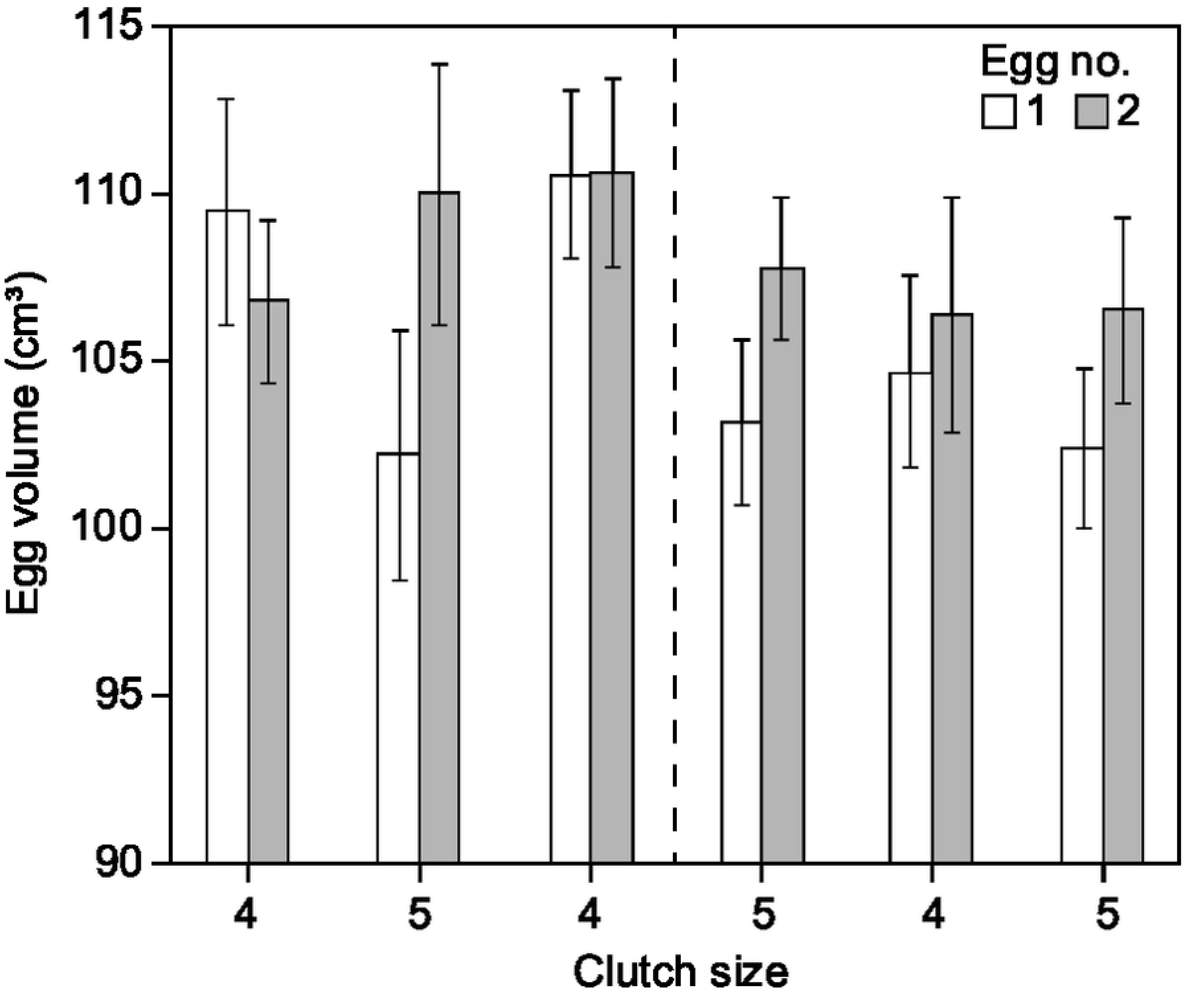
Average egg volume (±SE: standard error)) for egg number 1 and 2 in the laying sequence for individual female eiders that made two clutch size changes: from 4 to 5 to 4 eggs (N = 5) and from 5 to 4 to 5 eggs (N = 6) between years.

## Discussion

Our results confirmed a general intraclutch increase in egg size from the first to the second egg, with declining size in subsequent eggs, as commonly described in eiders and in several other Anatidae species [4, 7, 9, 47, 48]. However, we also found a consistent pattern of intraclutch egg-size variation depending on clutch size for the most commonly laid clutches of 4 and 5 eggs. In 4-egg clutches, the first egg was significantly larger and the second egg smaller than in 5-egg clutches (although just insignificant). This population level pattern in egg-size with clutch size was also found among individual females both when they increased and decreased their clutch size between seasons. As clutch size was significantly related to female body condition, our data suggest that intraclutch egg-size variation is an adaptive strategy adopted by individual females in response to their own pre-laying body condition.

### Clutch size variation

Female eider body condition prior to breeding is widely acknowledged as a major determinant of clutch size [23, 49, 50]. However, clutch size is also affected by female age and timing of laying [51, 52], which often are interrelated, with young birds laying later than older birds [53]. In this study, nest surveys were always performed in the early hatching period, and colony average laying date varied by only five days throughout the study period [40], suggesting no seasonal effect of timing on clutch sizes. Likewise, restricting the analysis to females caught on nests minimised the possible inclusion of late breeding young birds [cf. 54], as well as potential re-nesters. We therefore believe that the variation in average clutch size between seasons mainly reflects environmental stochasticity affecting conditions for breeding females.

### Intraclutch egg-size variation

Overall, the present data reflects the general pattern of intraclutch egg-size variation reported in many waterfowl species as well as in other eider populations [e.g., 7, 9, 11]. However, we show, for the first time, that egg size was significantly related to laying order (in 2-6 egg clutches), and the pattern of egg size variation with laying order was significantly affected by clutch size (in 2-6 egg clutches). This suggests that eider females produce eggs within a specific predetermined pattern of intraclutch size variation depending on ultimate clutch size. Since this pattern alternated in individual females when both increasing and decreasing clutch size between seasons (in relation to contemporary body mass), our results strongly support the interpretation that eider females actively adapt clutch and egg size as part of their reproductive strategy, following some pre-programmed clutch/egg size “formats”.

Overall, the absolute difference in the size of the first and second eggs in 4 and 5-egg clutches was ∼1.4 cm^3^, corresponding to approx. 1.2% of the average egg size of the first two eggs. This relatively small change in the first laid egg has not been commented in previous studies, even though matching patterns showing a larger first egg in 4-egg clutches compared to 5-egg clutches, have been reported in other eider populations (Figure 7 in [7]; Table 1 in [9], see also [10].

### Implications of changing egg size

Egg size is recognised as a major determinant of hatchling success, and increases in egg size directly affects the size of ducklings within and between clutches, and hence may have significant fitness value [55, 56,]. Reproductive success (in terms of ducklings leaving the nests and duckling survival) increases with clutch size [57–59]. Hence, increasing the size of the first laid egg in 4-egg clutches, has the potential to partly compensate for overall reduced reproductive success compared to a higher success in 5 and 6 egg clutches. Thus, when laying a 4-egg clutch, females produce two large eggs, which are almost comparable in size of the two largest eggs in a 6 egg clutch and comparable to the second largest egg in a 5-egg clutch (cf. Fig. 2). Hence, instead of producing only one large egg, if the size of the first egg was similar to that of 5 and 6 egg clutches, females laying 4-egg clutches lay two large eggs and hatch two relatively large ducklings, with the likely fitness benefits associated. In geese and large waterfowl, greater size at hatching is reported to benefit individuals by a stronger competitive ability for optimal brooding, more efficient foraging skills, stronger and quicker responses to predator warning by the female and/or a more efficient physiological/immunological state. All traits that probably contribute to higher survival [cf. 28, 60].

The advantage of laying two relatively large eggs may however, not only be restricted to the individual ducklings within a clutch of 4 eggs. As eiders frequently engage in brood amalgamation, forming so-called crèches [61, 62], larger ducklings may benefit from their size advantage at the multi-brood/cohort level when grouped with other ducklings from one or more broods, or abandoned by the female into crèches [but see 63]. In such a cohort ‘size-scenario’, the present data (cf. Fig. 2) shows that among the largest 20% of all ducklings in an average cohort, almost half (47.6%) originate from 4-egg clutches (from the first and second egg), 48.6% originate from 5-egg clutches (from the second and third egg), and 3.8% from 6 egg clutches (from the second and third egg). Without the recorded size increase in the first egg (compared to the size of first egg of 5 and 6 egg clutches), 4-egg clutches would only represent 31.2% of the largest ducklings in a cohort, while 63.8% and 4.9% will be from 5 and 6 egg clutches respectively. Hence, in terms of simple size, the egg size pattern of 4-egg females may be of adaptive advantage both in terms of reproductive success (duckling survival probability), as the relative duckling size within cohorts determine survival and recruitment [26], and by allowing females in a suboptimal body condition to reduce clutch size and/or abandon ducklings, to safeguard own survival [cf. 64].

### Compliance with general hypothesis

Most studies of intraclutch egg size in eiders and other precocial waterfowl show variation consistent with the adaptive resource distribution hypothesis, stating that females allocate most resources to eggs with a higher probability of survival, which reflects their position in the laying sequence [4, 8, 9, 11, 65, see also 36]. In eiders, highest energy investment, assessed here by larger egg size, is allocated to the first 3 eggs in the laying sequence. Of these, pre-incubation failure (mainly predation) of unattended first laid eggs are much higher than for subsequent eggs [66, 67], whereas higher survival in subsequent eggs are attributed to increased nest attendance and incubation initiation, which starts after the laying of the second or third eggs [10, 66, 67]. Hence, increasing the size of the first laid egg when laying 4 eggs, as found in the present study, seems a highly risky investment, if not accompanied by an earlier nest attendance. We did not investigate the timing of nest attendance and incubation initiation in the present study, but Hanssen et al. [10] showed that eiders laying 4-egg clutches start incubation on average one day earlier in the laying sequence than those laying 5-egg clutches. They also showed that an earlier start to incubation was related to poor female body condition, but they did not relate advanced incubation to a larger size of the first laid egg in 4-egg clutches, even though their data showed such a pattern (see Hanssen et al. [10]). In snow geese, Williams et al. [32] reported that larger first laid eggs had a higher survival prior to incubation start than smaller first laid eggs. They did not consider timing of the start of incubation in their study, but their results also indicated that female snow geese laying large first eggs in a clutch were capable of increasing survival of these eggs.

In theory, adaptive changes in egg size when laying a suboptimal clutch size, should occur in eggs of intermediate size and not in the smallest or largest eggs. Hence, in both 4, 5 (and 6) egg clutches, expected candidates for adaptive changes in size would be egg number one and three. The present analyses showed a size increase in the relatively large first laid egg, in line with our prediction, but this result somehow conflicts with the resource distribution hypothesis, as first laid eggs generally are reported to have a high failure rate (see above).

Although an earlier incubation start may represent an adaptation to increase first egg success in 4-egg clutches [cf. 10], a size increase in the third laid egg would have been more in line with the resource distribution hypothesis, with largest investment in the two central eggs. However, increasing the size of the third egg would probably postpone incubation initiation and further increase pre-incubation failure (predation risk) of first laid eggs, as incubation initiation may not commence prior to laying the largest egg within a clutch, without inducing a hatching or developmental asynchrony in the embryos [cf. 68]. Alternatively, the laying of two large consecutive first eggs of comparable size may facilitate an earlier incubation initiation, as females may increase nest attendance or start incubation after laying the first of these similar sized eggs, without jeopardizing synchronous development and hatching. If so, the laying of similar sized first and second egg in 4-egg clutches may explain the one day earlier incubation initiation in these clutch sizes [cf. 10]. Likewise, similar egg size in 6 egg clutches, where the second and third egg are of comparable size, may reflect an adaptation to advance incubation start from the third to the second egg. Although no data exist on the time of incubation start in 6 egg clutches, and sufficient data may be hard to obtain, an advanced incubation start, comparable to that for 5-egg clutches, could explain how reproductive success, expressed as successfully fledged ducklings, generally increases with clutch size [57, 58].

In conclusion, our study shows that individual female eiders have a specific within-clutch egg-size strategy, dependent on clutch size. Based on sequential histories of specific females, where individual females laid larger first eggs in a 4-egg clutch when in a poor condition, and laid smaller first eggs in a 5-egg clutch when in better condition, we argue, that this change is a functional adaptation, and that this adaptation may increase reproductive output, as the size increase in first egg results in two ducklings (rather than one) that benefit from large size, in both single or amalgamated broods. We suggest that the increased investment in first laid eggs, which normally suffer from a high predation rate is, at least partly, compensated by an earlier start of nest attendance and incubation, as other studies have reported advanced incubation start in females in poor condition laying 4-egg clutches. Hence, our results suggest that eiders possess a finely tuned egg and clutch size strategy related to their pre-laying body condition, by which individuals can adjust current reproduction in a varying environment.

## Acknowledgements

We are grateful to all the people who helped in catching eiders and to measure egg size during all years, especially Jens Peder Hounisen, Ebbe B. Hansen and the late Henning Noer. Niels Adamsen is sincerely thanked for supporting the logistics on the island of Saltholm, and the Saltholm Ejerlaug and Danish Forest and Nature Agency for providing permission to access the island. Data collection was part of a general investigation commissioned by the A/S Øresundsbrokonsortiet, who are thanked for permission to use data. Finally, Anthony D. Fox and Kevin K. Clausen are thanked for commenting on an earlier version of this manuscript.

